# A Large-scale Comparison of Cortical and Subcortical Structural Segmentation Methods in Alzheimer’s Disease: a Statistical Approach

**DOI:** 10.1101/2020.08.18.256321

**Authors:** Jafar Zamani, Ali Sadr, Amir-Homayoun Javadi

## Abstract

**Background:** Alzheimer’s disease (AD) is a neurodegenerative disease that leads to anatomical atrophy, as evidenced by magnetic resonance imaging (MRI). Automated segmentation methods are developed to help with the segmentation of different brain areas. However, their reliability has yet to be fully investigated. To have a more comprehensive understanding of the distribution of changes in AD, as well as investigating the reliability of different segmentation methods, in this study we compared volumes of cortical and subcortical brain segments, using automated segmentation methods in more than 60 areas between AD and healthy controls (HC).

**Methods:** A total of 44 MRI images (22 AD and 22 HC, 50% females) were taken from the minimal interval resonance imaging in Alzheimer’s disease (MIRIAD) dataset. HIPS, volBrain, CAT and BrainSuite segmentation methods were used for the subfields of hippocampus, and the rest of the brain.

**Results:** While HIPS, volBrain and CAT showed strong conformity with the past literature, BrainSuite misclassified several brain areas. Additionally, the volume of the brain areas that successfully discriminated between AD and HC showed a correlation with mini mental state examination (MMSE) scores. The two methods of volBrain and CAT showed a very strong correlation. These two methods, however, did not correlate with BrainSuite.

**Conclusion:** Our results showed that automated segmentation methods HIPS, volBrain and CAT can be used in the classification of AD and HC. This is an indication that such methods can be used to inform researchers and clinicians of underlying mechanisms and progression of AD.

## 1 Introduction

Alzheimer’s disease (AD) is a devasting neurodegenerative disease, contributing to 60-70% of dementia cases ^1^. Currently there are around 50 million people with dementia worldwide. In 2015, the total global societal cost of dementia was estimated to be US$ 818 billion ^2,3^ Mainly due to increased life-expectancy, the total number of people with dementia is projected to reach 82 million (64% increase) in 2030, with between 49 and 57 million of these cases being AD ^4^ Whilst several drugs are available to mitigate the symptoms in some cases, no treatments are available that prevent progression from the relatively late stage at which the disease is diagnosed ^5^.

AD is characterised by two main pathological hallmarks: extracellular amyloid deposits, composed of insoluble amyloid beta (Aβ) protein, and intra-neuronal neurofibrillary tangles (NFTs), containing hyperphosphorylated tau protein ^6^. AD is also characterised by a significant loss of neurons and synapses, resulting in brain shrinkage and atrophy ^7,8^. Structural changes have been shown to be one of the earliest biomarkers that can be used in the diagnosis of AD ^9–11^. Much effort has been devoted to find patterns of changes in the structure of different brain areas that can be reliably used for diagnosis of AD ^12^

Earlier investigations relied mostly on manual segmentation of brain areas requiring a great deal of expertise and time ^13–16^. Therefore, the majority of the focus has been devoted to changes in the hippocampus due to its distinct structure ^17^ It has been shown that a loss in hippocampal volume can be an indication of AD ^18,19^ Further investigations have looked at subfields of the hippocampus, showing a nonuniform rate of neuroplasticity due to their specialisation ^20–22^. For example, it has been shown that NFT begin in the medial temporal region and exhibit a characteristic distribution pattern across subfields, starting in the CA1 and later spreading to subiculum, CA2, CA3 and CA4/Dentate Gyrus ^23–27^.

With the development of semi- and fully-automated segmentation methods, however, it has now become easier and faster to segment not only the hippocampal area, but also other brain areas ^28–32^. HIPpocampus subfield Segmentation (HIPS) ^33^, volBrain ^34^, Computational Anatomy Toolbox (CAT) ^35,36^, BrainSuite ^37,38^ and FreeSurfer ^39^ are some of the commonly used semi- and fully-automated methods. These methods, however, are yet under development ^40,41^. For example, CA1 segmentation in the FreeSurfer v5.3 was partially included in the subiculum ^18^ potentially explaining why the CA1 field was reported to be insensitive to AD pathology in some ^42,43^ but not all ^44,45^. Similar findings have recently raised questions and concerns regarding the accuracy and consistency of these methods ^46–49^ Therefore, it is important to investigate the accuracy of these methods further ^50^.

Benefiting from the computational power of automated methods, analysis of a large number of brain images has become more feasible. Large datasets of brain scans such as Minimal Interval Resonance Imaging in Alzheimer’s Disease (MIRIAD), a public database of Alzheimer’s magnetic resonance imaging (MRI) ^51^, offer a great opportunity to have a more comprehensive approach to the underlying mechanism and progression of AD ^52–56^. It also facilitates multisite studies to form a more accurate understanding of the disease ^57,58^.

Mini mental state examination (MMSE) is one of the commonly accepted measurements of cognitive ability, in particular in clinical settings ^59–61^. This measure has been widely used in classification of AD. For example, MIRIAD classifies participants with score between 12 and 26/30 as AD and those higher than 26/30 as healthy control. There is huge body of literature showing correlation between MMSE score and brain atrophy ^62–64^

The aim of this study was to investigate the reliability of four automated segmentation methods of volBrain, CAT and BrainSuite for segmentation of the whole brain, and HIPS for segmentation of subfields of hippocampus, which belongs to the same analysis tool as volBrain. We used images belonging to MIRIAD. Correlation of the volume of each brain area with MMSE scores are also investigated. To investigate the reliability of the three methods volBrain, CAT and BrainSuite, the correlation of their common brain areas is also reported.

## 2 Material and Methods

### 2.1 Subjects

We used 44 images (22 AD and 22 HC) taken from the Minimal Interval Resonance Imaging in Alzheimer’s Disease (MIRIAD) dataset ^51^. Table 1 shows a summary of the descriptives of the participants.

**Table 1.** Comparison between Alzheimer’s disease (AD) and healthy controls (HC).

| **Measure** | **Alzheimer's Disease** | **Healthy Control** | ***p*** |
| --- | --- | --- | --- |
| n | 22 | 22 | - |
| Female (n [%]) | 11 [50] | 11 [50] | - |
| Age (mean [SD]) | 70.64 [5.55] | 70.61 [7.54] | 0.916 |
| MMSE (mean [SD]; min-max) | 17.272 [5.461]; 5-26 | 29.087 [1.083]; 27-30 | < 0.001 |

### 2.2 Magnetic Resonance Imaging (MRI)

Data was extracted from MIRIAD database. All subjects underwent MRI scanning on a 1.5 T Signa scanner (GE Medical Systems, Milwaukee, WI, USA). T1-weighted volumetric images were obtained using an inversion recovery prepared fast spoiled gradient echo sequence with acquisition parameters time to repetition = 15 ms, time to echo = 5.4 ms, flip angle = 15°, TI = 650 ms, a 24-cm field of view and a 256 × 256 matrix, to provide 124 contiguous 1.5-mm thick slices in the coronal plane (voxels 0.9735 × 0.9735 × 1.5 mm^3^) ^51^.

### 2.3 Methods

#### HIPS and volBrain

The volumes of Cerebrospinal fluid (CSF), white matter (WM), grey matter (GM), brain hemispheres, cerebellum and brainstem were obtained using volBrain pipeline ^34^. This method is based on an advanced pipeline providing automatic segmentation of different brain structures from T1 weighted MRI, Figure 1. The preprocessing is based on the following procedure: (1) a denoising step with an adaptive non-local mean filter, (2) an affine registration in the Montreal Neurological Institute (MNI) space, (3) a correction of the image inhomogeneities, and (4) an intensity normalisation. (5) Afterwards, MRI images are segmented in the MNI space using non-local patch-based multi-atlas method. Images were corrected for intensity inhomogeneity using the N4 algorithm ^65^, and the images were segmented into brain/non-brain using a semi-automated technique (MIDAS). The Non-Local Means filter ^66^ was applied to each pixel of the image by computing a weighted average of surrounding pixels using a robust similarity measure that takes into account the neighbouring pixels surrounding the pixel being compared. This segmentation method is based on the idea of non-local patch-based label fusion technique, where patches of the brain image to be segmented are compared with those of the training library, looking for similar patterns within a defined search volume to assign the proper label ^67,68^. HIPS and volBrain are used for segmentation of the hippocampus subfields and the rest of the brain, respectively ^33^.

**Figure 1.**
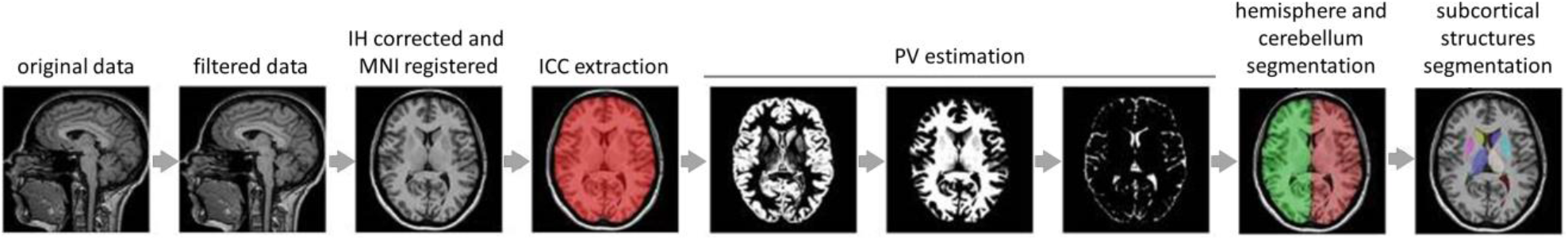
Processing pipeline for volBrain and HIPS adapted from Manjón and Coupé (2016) ^34^ under the terms of the Creative Commons Attribution License (CC BY).

#### CAT

Computational Anatomy Toolbox (CAT) is a powerful package for brain T1-MRI data segmentation, Figure 2. It is a voxel base estimation method ^69^. The CAT preprocessing steps are as follows: (1) spatial registration to a template, (2) tissue segmentation into grey, white matter and CSF, and (3) bias correction of intensity non-uniformities. (4) Finally, segments are extracted by scaling the amount of volume changes based on spatial registration, so that the total volume of grey matter in the modulated image remains the same as the original image. For correction of the orientation and size of the brain, non-linear registration methods are applied to the image ^70^. Projection-based thickness (PBT) method is used for calculation of the cortical thickness and central surface ^70,71^. Spatial-adaptive Non-Local Means (SANLM) and classical Markov Random Field (MRF) were used for image Denoising ^72^. Adaptive Maximum a Posterior (AMAP) method was used for segmentation ^69^

**Figure 2.**
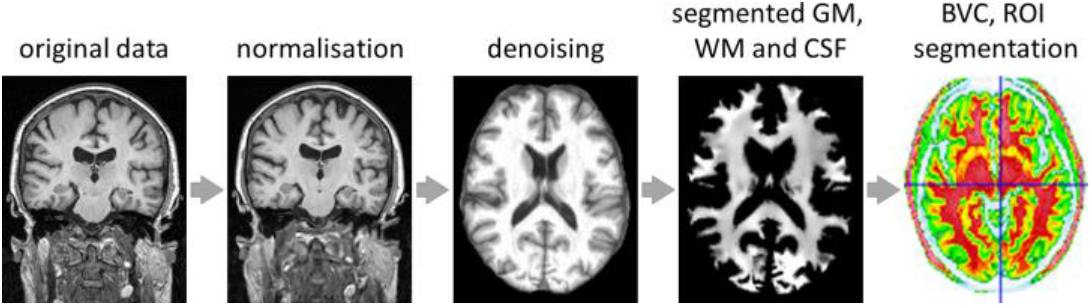
Processing pipeline for CAT

#### BrainSuite

BrainSuite is an open source software tool that enables largely automated cortical surface extraction from MRI of the brain, Figure 3. BrainSuite includes automatic cortical surface extraction, bias field correlation, cerebrum labelling, and surface generation features. Also, this toolbox is used in tractography and connectivity matrix calculation in diffusion imaging data ^37^.

**Figure 3.**
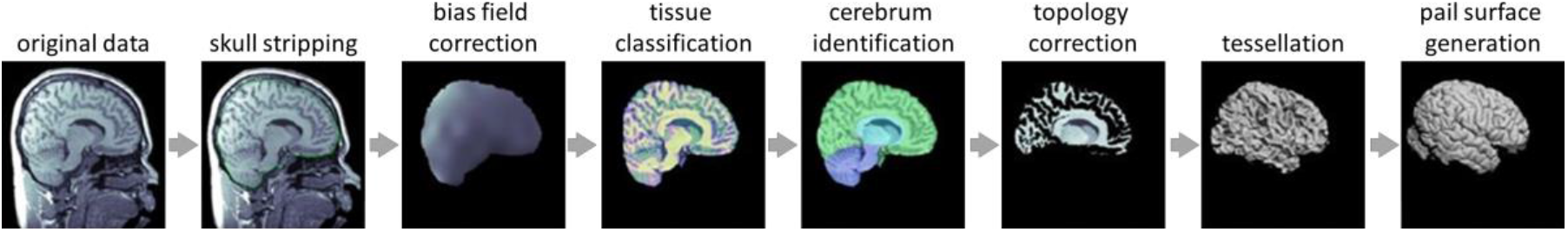
Processing pipeline for BrainSuite

### 2.4 Statistical Analysis

Independent-sample t-tests are run to compare the volume of different brain areas between the AD and HC groups for volBrain, CAT and BrainSuite for the whole brain, and HIPS for the hippocampus subfields. Bivariate-correlation analyses are also run to investigate the relationship between volume and MMSE scores for all four segmentation methods. Correlational analyses are run between the common brain areas in volBrain, CAT and BrainSuite to investigate the relationship between the three methods. Bonferroni correction is applied to account for multiple comparison.

## 3 Results

Using three automatic segmentation methods CAT, volBrain and BrainSuite, we segmented the whole brain, and using HIPS we segmented the hippocampus. Using independent-sample t-tests we compared the volumetric data for AD and HC for each segment. Figures 3–6 show sample output images for one AD patient and one HC participant. Furthermore, we investigated the correlation of volumetric data with MMSE scores in both groups.

**Figure 4.**
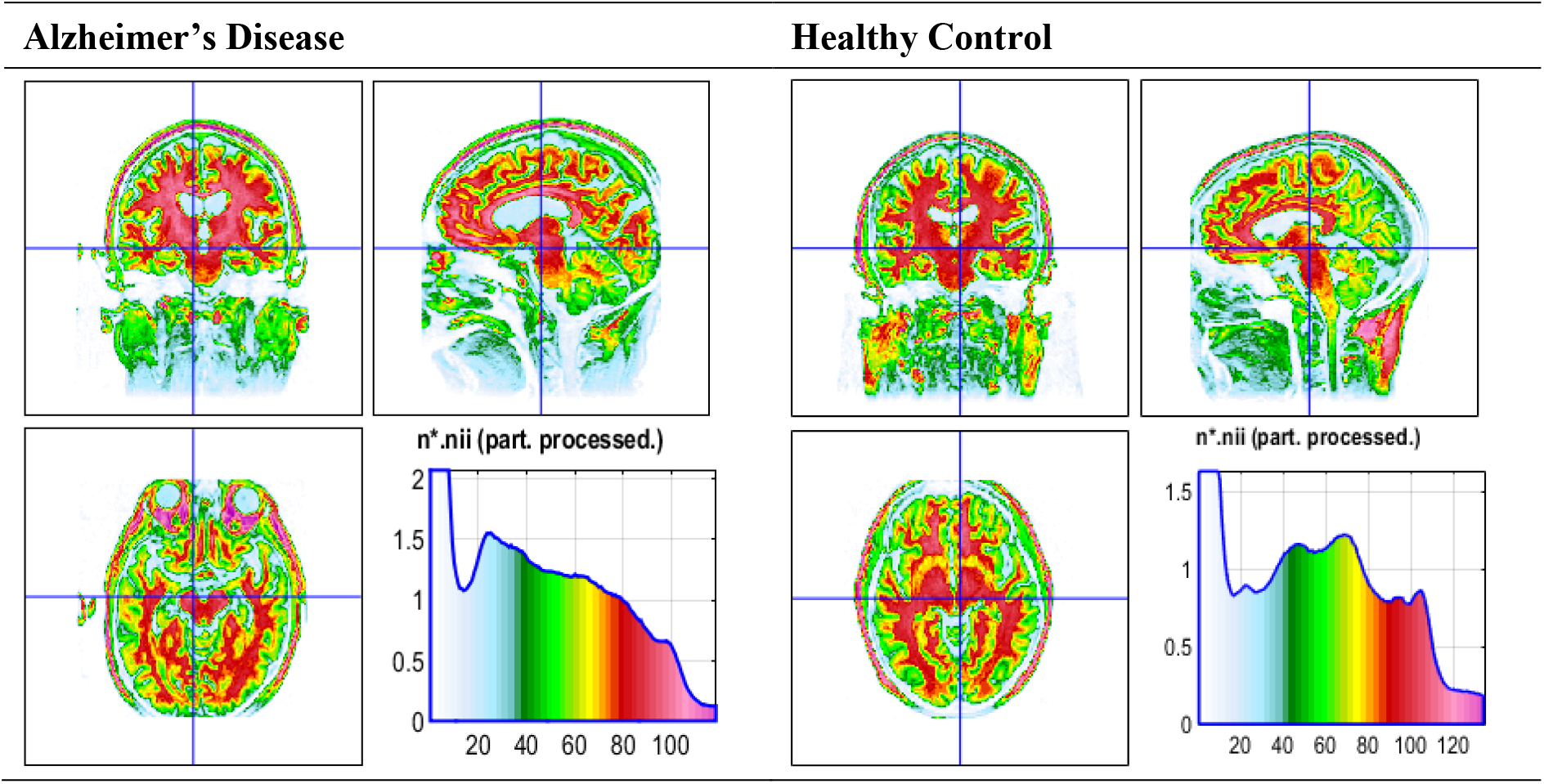
Subcortical structures in an AD patient and a HC participant using CAT segmentation method. The histograms show the volume of each brain area

**Figure 5.**
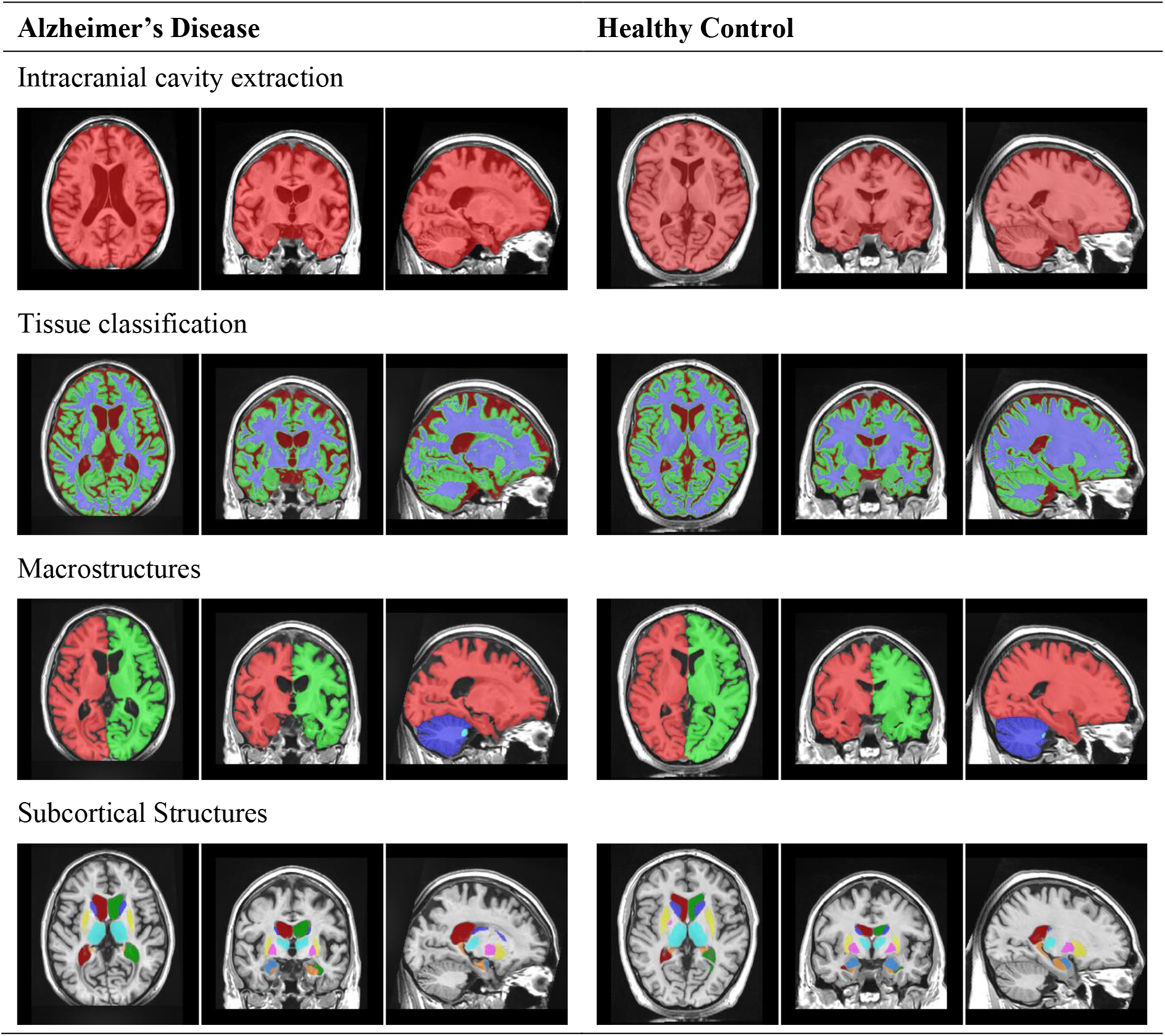
Cerebellum MRI-T1 brain segmentation in an AD patient and a HC participant using volBrain segmentation method.

**Figure 6.**
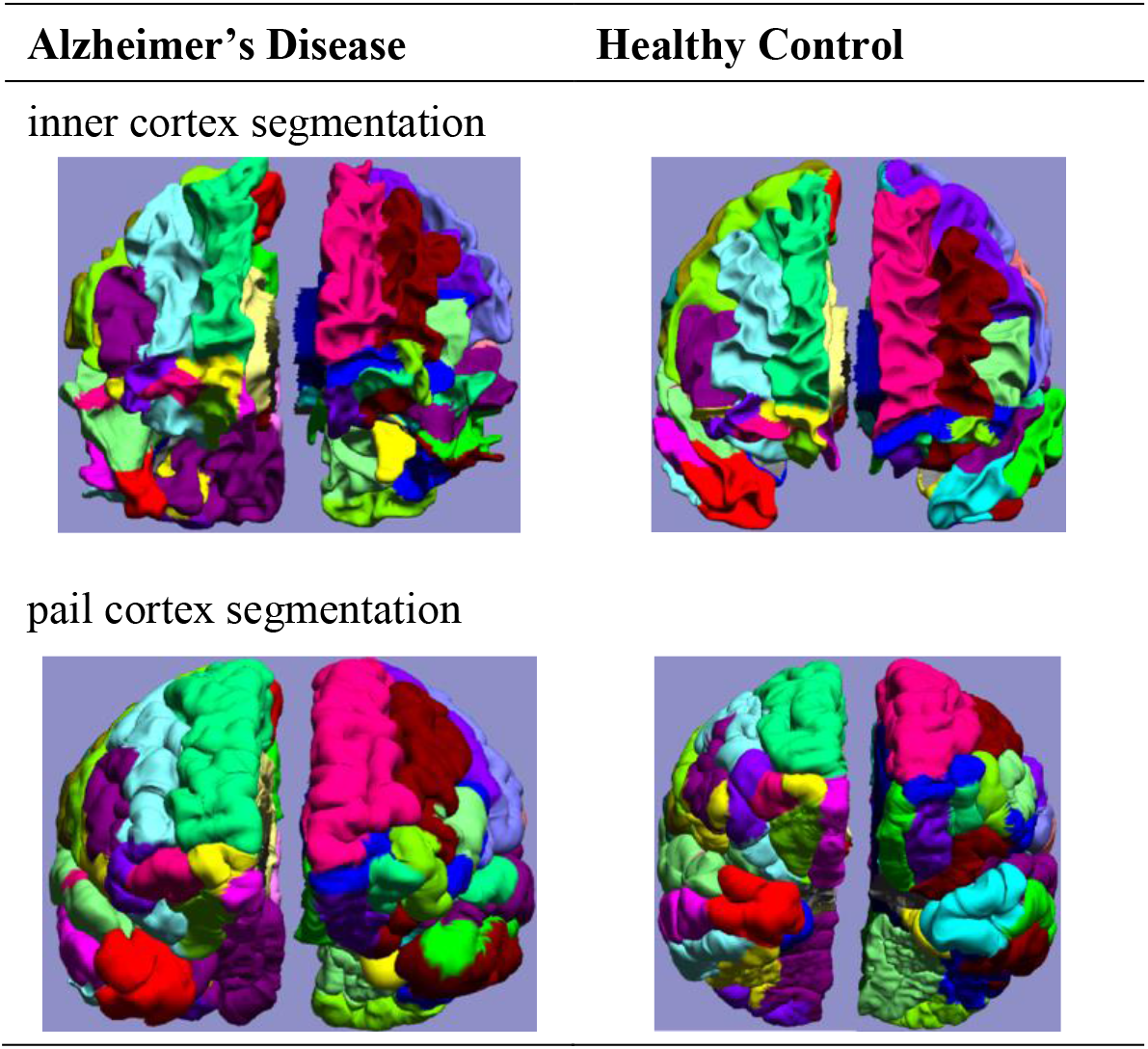
Cerebellum MRI-T1 brain segmentation in an AD patient and a HC participant using BrainSuite segmentation method.

**Figure 7.**
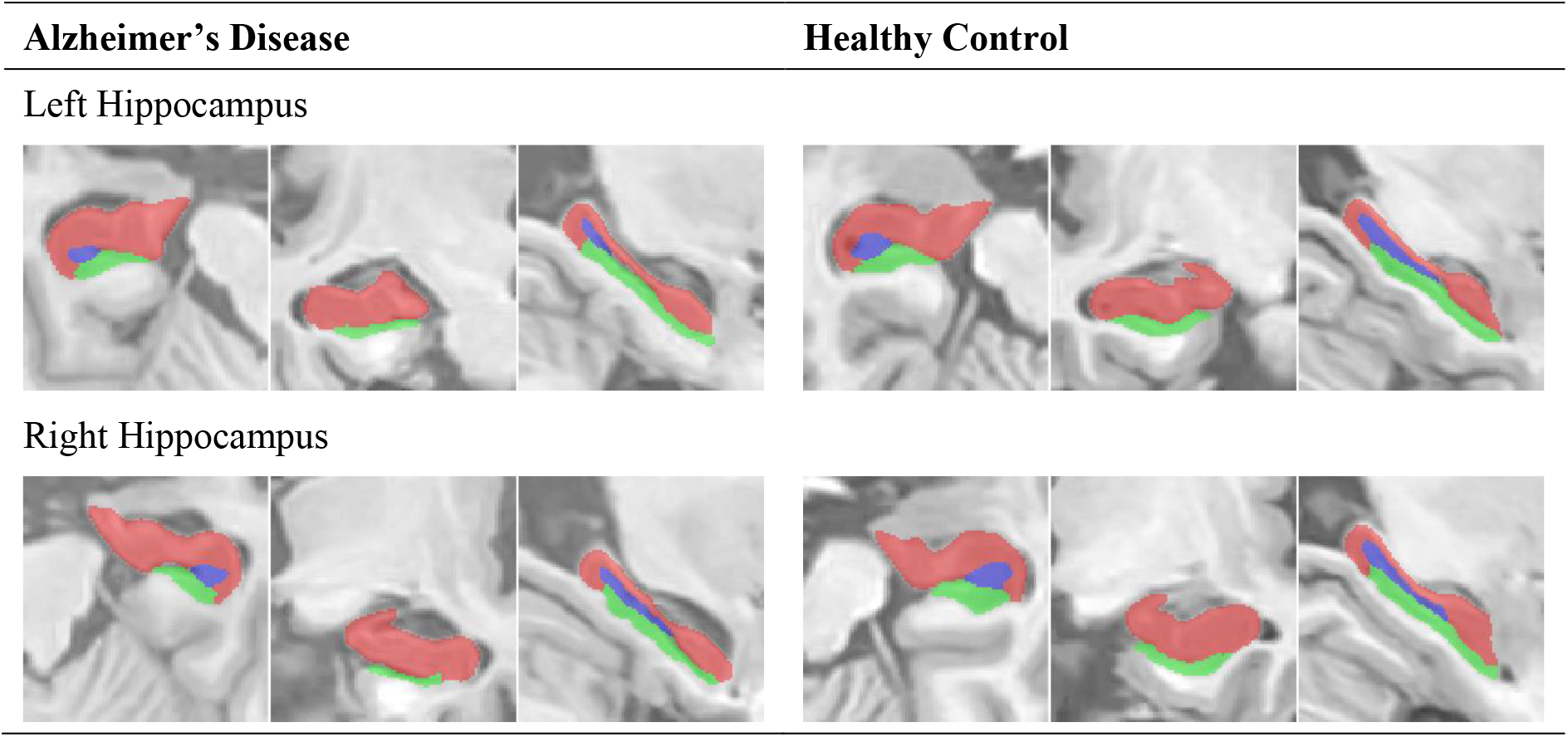
Left and right hippocampus subfield segmentation in an AD patient and a HC participant using HIPS segmentation method.

CAT segmentation method returned data for 63 distinct brain areas. This method highlighted many brain areas that are significantly different between the two groups, Table 2. In particular fusiform gyrus, parahippocampal gyrus, hippocampus, entorhinal cortex, amygdala, temporal gyri, thalamus, nucleus accumbens, insula, caudate and precuneus were significantly different. Importantly, the size of all these brain areas showed a strong correlation with MMSE scores. For further details see supplementary figures 1-3.

**Table 2.** Summary of the independent-sample t-tests comparing volumetric data between the participants with Alzheimer’s disease and healthy controls and the correlation of the data with MMSE scores using **CAT** method.

| Brain Area | Comparison between groups |  |  | Correlation with MMSE |  |
| --- | --- | --- | --- | --- | --- |
|  | *t* | *p*↑ | *d* | *r*<sup>2</sup> | *p* |
| Fusiform Gyrus | 10.793 | < 0.000001* | 3.413 | 0.723 | < 0.000001* |
| Parahippocampus Gyrus | 9.936 | < 0.000001* | 3.142 | 0.536 | < 0.000001* |
| Hippocampus | 9.753 | < 0.000001* | 3.084 | 0.460 | 0.000001* |
| Entorhinal Cortex | 9.717 | < 0.000001* | 3.073 | 0.476 | < 0.000001* |
| Amygdala | 9.043 | < 0.000001* | 2.860 | 0.445 | 0.000001* |
| Inferior Temporal Gyrus | 8.939 | < 0.000001* | 2.827 | 0.653 | < 0.000001* |
| Middle Temporal Gyrus | 7.632 | < 0.000001* | 2.413 | 0.619 | < 0.000001* |
| Temporal Pole | 7.185 | < 0.000001* | 2.272 | 0.491 | < 0.000001* |
| Basal Forebrain | 6.658 | < 0.000001* | 2.105 | 0.453 | 0.000001* |
| Thalamus | 6.344 | < 0.000001* | 2.006 | 0.471 | 0.000001* |
| Angular Gyrus | 5.808 | 0.000001* | 1.837 | 0.507 | < 0.000001* |
| Accumbens | 5.275 | 0.000005* | 1.668 | 0.289 | 0.000236* |
| Inferior Occipital Gyrus | 5.228 | 0.000006* | 1.653 | 0.527 | < 0.000001* |
| Superior Temporal Gyrus | 5.186 | 0.000007* | 1.640 | 0.513 | < 0.000001* |
| Supramarginal Gyrus | 5.101 | 0.000009* | 1.613 | 0.498 | < 0.000001* |
| Anterior Insula | 4.955 | 0.000014* | 1.567 | 0.346 | 0.000043* |
| Occipital Fusiform Gyrus | 4.519 | 0.000054* | 1.429 | 0.472 | 0.000001* |
| Middle Occipital Gyrus | 4.515 | 0.000055* | 1.428 | 0.417 | 0.000004* |
| Posterior Insula | 4.447 | 0.000068* | 1.406 | 0.338 | 0.000054* |
| Planum Polare | 4.232 | 0.000131* | 1.338 | 0.349 | 0.000038* |
| Anterior Cingulate Gyrus | 4.225 | 0.000134* | 1.336 | 0.362 | 0.000025* |
| Superior Parietal Lobule | 4.131 | 0.000179* | 1.306 | 0.423 | 0.000003* |
| Caudate | 3.959 | 0.000301* | 1.252 | 0.234 | 0.001155 |
| Subcallosal Area | 3.847 | 0.000420* | 1.217 | 0.299 | 0.000180* |
| Middle Frontal Gyrus | 3.796 | 0.000490* | 1.200 | 0.328 | 0.000073* |
| Medial Orbital Gyrus | 3.739 | 0.000579* | 1.182 | 0.278 | 0.000331* |
| Inferior Frontal Gyrus | 3.691 | 0.000666* | 1.167 | 0.373 | 0.000017* |
| Precuneus | 3.633 | 0.000788* | 1.149 | 0.349 | 0.000038* |
| Superior Medial Frontal Gyrus | 3.543 | 0.001023 | 1.120 | 0.312 | 0.000120* |
| Putamen | 3.525 | 0.001077 | 1.115 | 0.203 | 0.002822 |
| Temporal | 3.489 | 0.001195 | 1.103 | 0.340 | 0.000051* |
| Anterior Orbital Gyrus | 3.413 | 0.001484 | 1.079 | 0.244 | 0.000894 |
| Posterior Orbital Gyrus | 3.272 | 0.002203 | 1.035 | 0.255 | 0.000643* |
| Lingual Gyrus | 3.263 | 0.002262 | 1.032 | 0.304 | 0.000158* |
| Posterior Cingulate Gyrus | 3.226 | 0.002506 | 1.020 | 0.314 | 0.000117* |
| Central Operculum | 3.116 | 0.003388 | 0.985 | 0.372 | 0.000018* |
| Frontal Operculum | 2.999 | 0.004638 | 0.948 | 0.265 | 0.000479* |
| Supplementary Motor Cortex | 2.969 | 0.005032 | 0.939 | 0.310 | 0.000128* |
| Exterior Cerebellum | 2.934 | 0.005513 | 0.928 | 0.163 | 0.007986 |
| Superior Frontal Gyrus | 2.870 | 0.006523 | 0.908 | 0.199 | 0.003066 |
| Parietal Operculum | 2.864 | 0.006637 | 0.906 | 0.320 | 0.000095* |
| Middle Cingulate Gyrus | 2.734 | 0.009272 | 0.865 | 0.269 | 0.000432* |
| Gyrus Rectus | 2.660 | 0.011190 | 0.841 | 0.143 | 0.013603 |
| Optic Chiasm | 2.531 | 0.015401 | 0.800 | 0.066 | 0.101456 |
| Lateral Orbital Gyrus | 2.427 | 0.019841 | 0.767 | 0.158 | 0.009194 |
| Temporal Transverse Gyrus | 2.345 | 0.024056 | 0.742 | 0.189 | 0.003960 |
| Superior Occipital Gyrus | 2.290 | 0.027347 | 0.724 | 0.245 | 0.000849 |
| Medial Precentral Gyrus | 2.250 | 0.030015 | 0.712 | 0.196 | 0.003321 |
| Cuneus | 2.091 | 0.042968 | 0.661 | 0.187 | 0.004135 |
| Inferior Frontal Orbital Gyrus | 2.063 | 0.045679 | 0.652 | 0.123 | 0.022887 |
| Postcentral Gyrus | 1.687 | 0.099402 | 0.533 | 0.142 | 0.013747 |
| Frontal Pole | 1.603 | 0.116854 | 0.507 | 0.067 | 0.098513 |
| Occipital Pole | 1.596 | 0.118311 | 0.505 | 0.127 | 0.020199 |
| Inferior Frontal Angular Gyrus | 1.518 | 0.136864 | 0.480 | 0.134 | 0.017041 |
| Cerebellum White Matter | 1.319 | 0.194795 | 0.417 | 0.079 | 0.071875 |
| Precentral Gyrus | 1.196 | 0.238644 | 0.378 | 0.124 | 0.022441 |
| Medial Postcentral Gyrus | 1.061 | 0.294904 | 0.336 | 0.070 | 0.090174 |
| Brainstem | -0.875 | 0.386852 | 0.277 | 0.005 | 0.640175 |
| Cerebellar Vermal Lobules VI-VII | 0.699 | 0.488487 | 0.221 | 0.024 | 0.325448 |
| Cerebellar Vermal Lobules VIII-X | 0.617 | 0.541029 | 0.195 | 0.015 | 0.434774 |
| Cerebellar Vermal Lobules I-V | 0.218 | 0.828189 | 0.069 | 0.016 | 0.430517 |
| Globus Pallidus | -0.212 | 0.832833 | 0.067 | 0.001 | 0.860265 |
| Calcarine Cortex | 0.198 | 0.843964 | 0.063 | 0.013 | 0.473158 |

Notes: ↑ rows are sorted based on the $p$ values for the t-test; \* $p < 0.000793$ Bonferroni corrected for multiple comparison; $d$ represents Cohen's $d$ effect size; MMSE: mini mental state examination

volBrain segmentation method returned data for eight distinct brain areas. In particular the amygdala, hippocampus, nucleus accumbens, thalamus and caudate were significantly different between the two groups, Table 3. Again, the size of all these brain areas showed a strong correlation with MMSE scores. For further details see supplementary figures 4-6.

**Table 3.** Summary of the independent-sample t-tests comparing volumetric data between the participants with Alzheimer’s disease and healthy controls and the correlation of the data with MMSE scores using **volBrain** method.

| Brain Area | Comparison between groups |  |  | Correlation with MMSE |  |
| --- | --- | --- | --- | --- | --- |
|  | *t* | *p*↑ | *d* | *r*<sup>2</sup> | *p* |
| Amygdala | 10.217 | < 0.000001* | 3.231 | 0.428 | 0.000001* |
| Hippocampus | 6.58 | < 0.000001* | 2.081 | 0.256 | 0.000395* |
| Accumbens | 5.813 | 0.000001* | 1.838 | 0.339 | 0.000027* |
| Thalamus | 4.422 | 0.000065* | 1.398 | 0.317 | 0.000057* |
| Caudate | 4.149 | 0.000154* | 1.312 | 0.169 | 0.005091 |
| Cerebellum | 2.063 | 0.045216 | 0.652 | 0.094 | 0.041135 |
| Globus Pallidus | -1.103 | 0.276245 | 0.349 | 0.026 | 0.287427 |
| Putamen | 0.846 | 0.402030 | 0.268 | 0.019 | 0.366267 |

Notes: ↑ rows are sorted based on the *p* values for the t-test; \* *p* < 0.0038 Bonferroni corrected for multiple comparison; *d* represents Cohen's *d* effect size; MMSE: mini mental state examination;

BrainSuite segmentation method returned data for 50 distinct brain areas. In contrast to CAT and volBrain, this method highlighted only six brain areas that are significantly different between the two groups, Table 4. These brain areas included temporal gyri, third ventricle, supramarginal gyrus and angular gyrus. Similar to previous segmentation methods, all these brain areas showed strong correlation with MMSE scores. For further details see supplementary figures 7-9.

**Table 4.**
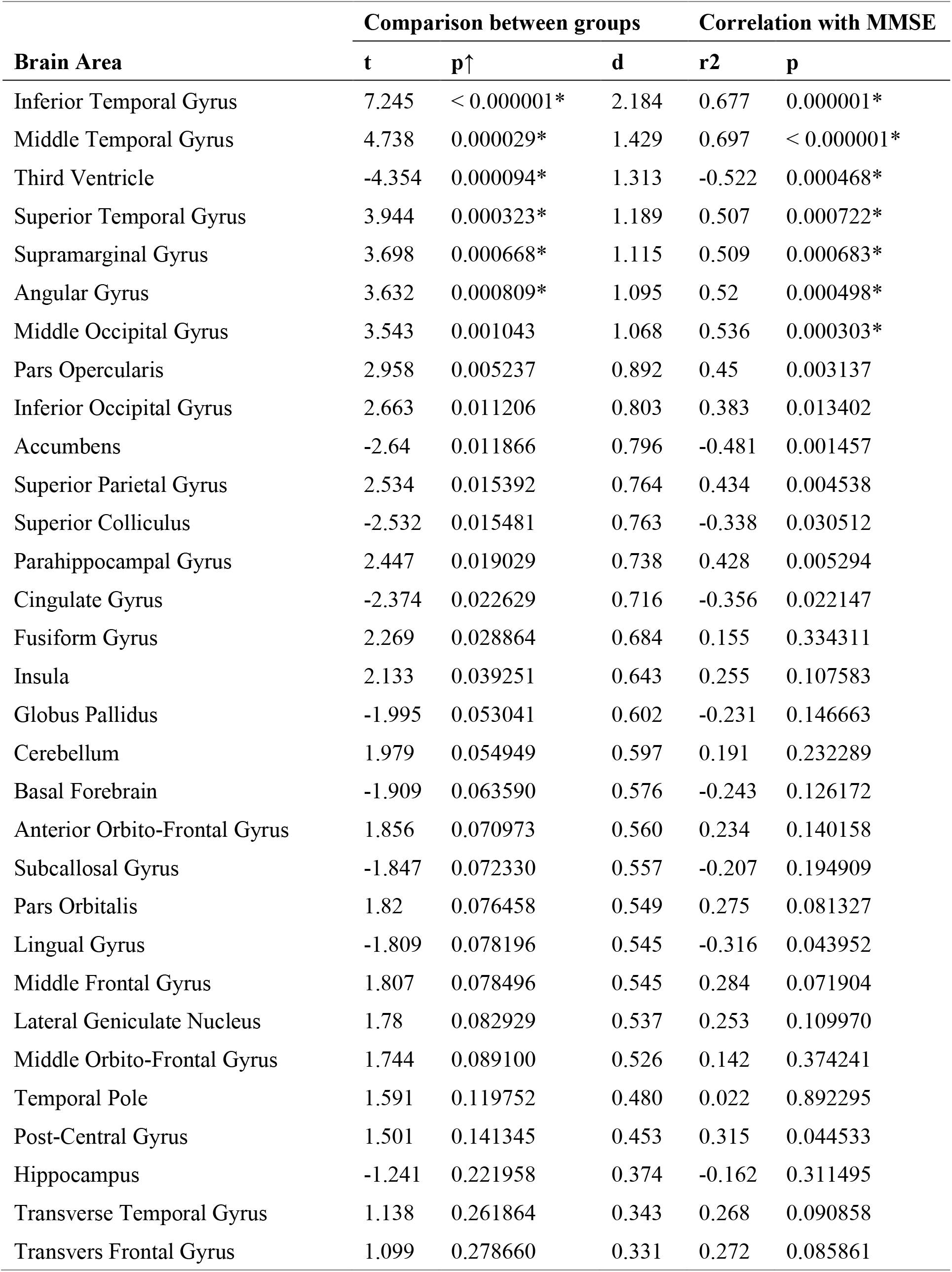
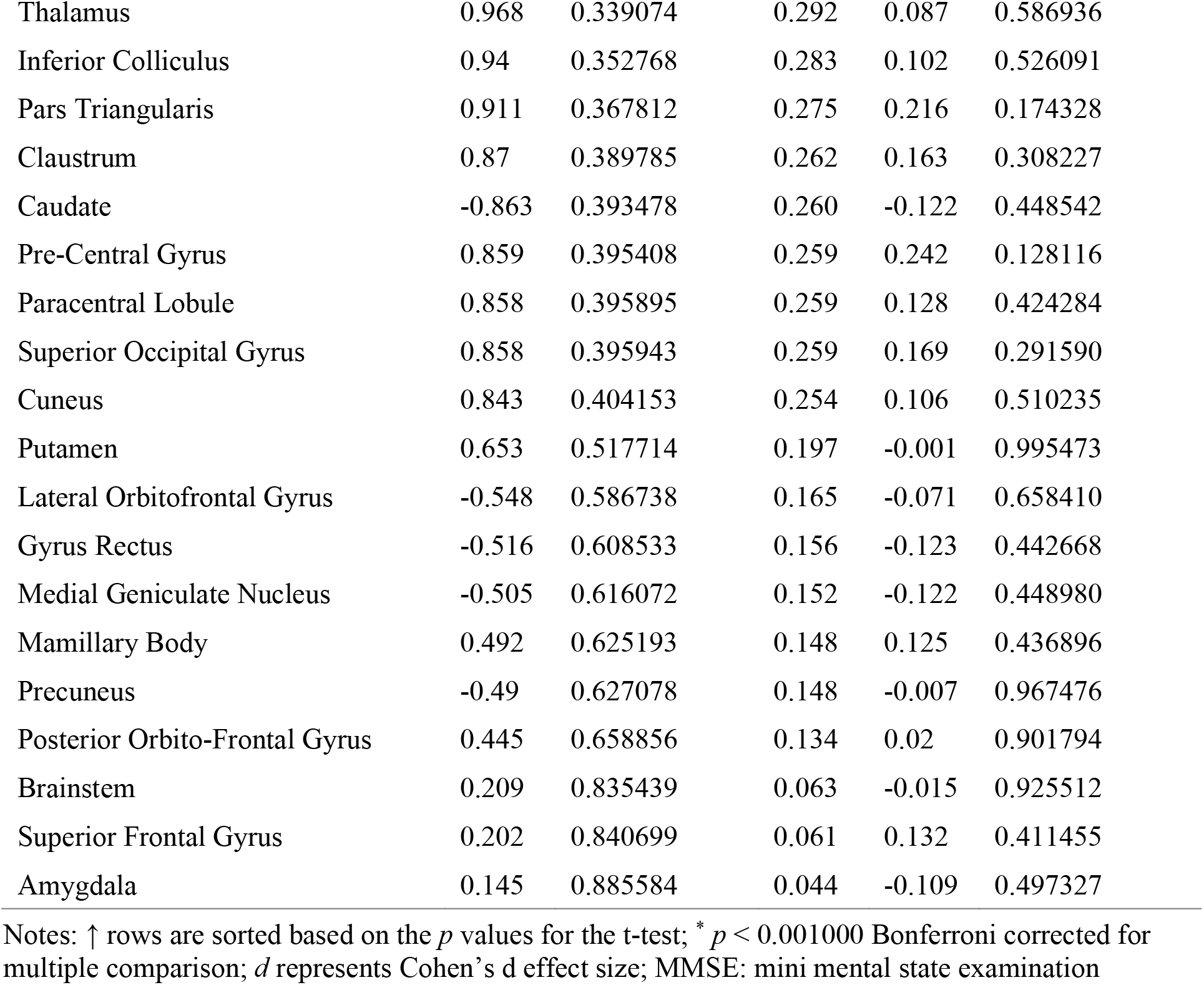
Summary of the independent-sample t-tests comparing volumetric data between the participants with Alzheimer’s disease and healthy controls and the correlation of the data with MMSE scores using **BrainSuite** segmentation method.

| Brain Area | Comparison between groups |  |  | Correlation with MMSE |  |
| --- | --- | --- | --- | --- | --- |
|  | t | p↑ | d | r2 | p |
| Inferior Temporal Gyrus | 7.245 | < 0.000001* | 2.184 | 0.677 | 0.000001* |
| Middle Temporal Gyrus | 4.738 | 0.000029* | 1.429 | 0.697 | < 0.000001* |
| Third Ventricle | -4.354 | 0.000094* | 1.313 | -0.522 | 0.000468* |
| Superior Temporal Gyrus | 3.944 | 0.000323* | 1.189 | 0.507 | 0.000722* |
| Supramarginal Gyrus | 3.698 | 0.000668* | 1.115 | 0.509 | 0.000683* |
| Angular Gyrus | 3.632 | 0.000809* | 1.095 | 0.52 | 0.000498* |
| Middle Occipital Gyrus | 3.543 | 0.001043 | 1.068 | 0.536 | 0.000303* |
| Pars Opercularis | 2.958 | 0.005237 | 0.892 | 0.45 | 0.003137 |
| Inferior Occipital Gyrus | 2.663 | 0.011206 | 0.803 | 0.383 | 0.013402 |
| Accumbens | -2.64 | 0.011866 | 0.796 | -0.481 | 0.001457 |
| Superior Parietal Gyrus | 2.534 | 0.015392 | 0.764 | 0.434 | 0.004538 |
| Superior Colliculus | -2.532 | 0.015481 | 0.763 | -0.338 | 0.030512 |
| Parahippocampal Gyrus | 2.447 | 0.019029 | 0.738 | 0.428 | 0.005294 |
| Cingulate Gyrus | -2.374 | 0.022629 | 0.716 | -0.356 | 0.022147 |
| Fusiform Gyrus | 2.269 | 0.028864 | 0.684 | 0.155 | 0.334311 |
| Insula | 2.133 | 0.039251 | 0.643 | 0.255 | 0.107583 |
| Globus Pallidus | -1.995 | 0.053041 | 0.602 | -0.231 | 0.146663 |
| Cerebellum | 1.979 | 0.054949 | 0.597 | 0.191 | 0.232289 |
| Basal Forebrain | -1.909 | 0.063590 | 0.576 | -0.243 | 0.126172 |
| Anterior Orbito-Frontal Gyrus | 1.856 | 0.070973 | 0.560 | 0.234 | 0.140158 |
| Subcallosal Gyrus | -1.847 | 0.072330 | 0.557 | -0.207 | 0.194909 |
| Pars Orbitalis | 1.82 | 0.076458 | 0.549 | 0.275 | 0.081327 |
| Lingual Gyrus | -1.809 | 0.078196 | 0.545 | -0.316 | 0.043952 |
| Middle Frontal Gyrus | 1.807 | 0.078496 | 0.545 | 0.284 | 0.071904 |
| Lateral Geniculate Nucleus | 1.78 | 0.082929 | 0.537 | 0.253 | 0.109970 |
| Middle Orbito-Frontal Gyrus | 1.744 | 0.089100 | 0.526 | 0.142 | 0.374241 |
| Temporal Pole | 1.591 | 0.119752 | 0.480 | 0.022 | 0.892295 |
| Post-Central Gyrus | 1.501 | 0.141345 | 0.453 | 0.315 | 0.044533 |
| Hippocampus | -1.241 | 0.221958 | 0.374 | -0.162 | 0.311495 |
| Transverse Temporal Gyrus | 1.138 | 0.261864 | 0.343 | 0.268 | 0.090858 |
| Transvers Frontal Gyrus | 1.099 | 0.278660 | 0.331 | 0.272 | 0.085861 |
| Thalamus | 0.968 | 0.339074 | 0.292 | 0.087 | 0.586936 |
| Inferior Colliculus | 0.94 | 0.352768 | 0.283 | 0.102 | 0.526091 |
| Pars Triangularis | 0.911 | 0.367812 | 0.275 | 0.216 | 0.174328 |
| Clastrum | 0.87 | 0.389785 | 0.262 | 0.163 | 0.308227 |
| Caudate | -0.863 | 0.393478 | 0.260 | -0.122 | 0.448542 |
| Pre-Central Gyrus | 0.859 | 0.395408 | 0.259 | 0.242 | 0.128116 |
| Paracentral Lobule | 0.858 | 0.395895 | 0.259 | 0.128 | 0.424284 |
| Superior Occipital Gyrus | 0.858 | 0.395943 | 0.259 | 0.169 | 0.291590 |
| Cuneus | 0.843 | 0.404153 | 0.254 | 0.106 | 0.510235 |
| Putamen | 0.653 | 0.517714 | 0.197 | -0.001 | 0.995473 |
| Lateral Orbitofrontal Gyrus | -0.548 | 0.586738 | 0.165 | -0.071 | 0.658410 |
| Gyrus Rectus | -0.516 | 0.608533 | 0.156 | -0.123 | 0.442668 |
| Medial Geniculate Nucleus | -0.505 | 0.616072 | 0.152 | -0.122 | 0.448980 |
| Mamillary Body | 0.492 | 0.625193 | 0.148 | 0.125 | 0.436896 |
| Precuneus | -0.49 | 0.627078 | 0.148 | -0.007 | 0.967476 |
| Posterior Orbito-Frontal Gyrus | 0.445 | 0.658856 | 0.134 | 0.02 | 0.901794 |
| Brainstem | 0.209 | 0.835439 | 0.063 | -0.015 | 0.925512 |
| Superior Frontal Gyrus | 0.202 | 0.840699 | 0.061 | 0.132 | 0.411455 |
| Amygdala | 0.145 | 0.885584 | 0.044 | -0.109 | 0.497327 |

Notes: ↑ rows are sorted based on the $p$ values for the t-test; \* $p < 0.001000$ Bonferroni corrected for multiple comparison; $d$ represents Cohen's $d$ effect size; MMSE: mini mental state examination

HIPS segmentation method returned data for the whole hippocampus and five of its subfields: CA1, CA2-CA3, CA4/Dentate Gyrus, Subiculum and strata radiatum/lacunosum/moleculare (SR-SL-SM). All these areas showed a significant difference between the two groups, Table 5. The size of hippocampus and all its subfields showed strong correlation with MMSE scores. For further details see supplementary figures 10-12.

**Table 5.** Summary of the independent-sample t-tests comparing volumetric data between the participants with Alzheimer’s disease and healthy controls and the correlation of the data with MMSE scores using **HIPS** segmentation method.

| Brain Area | Comparison between groups |  |  | Correlation with MMSE |  |
| --- | --- | --- | --- | --- | --- |
|  | *t* | *p*↑ | *d* | *r*<sup>2</sup> | *p* |
| SR-SL-SM | 8.990 | < 0.000001* | 2.843 | 0.393 | 0.000004* |
| Hippocampus | 8.619 | < 0.000001* | 2.726 | 0.388 | 0.000005* |
| CA4/Dentate Gyrus | 8.248 | < 0.000001* | 2.608 | 0.402 | 0.000003* |
| CA1 | 6.308 | < 0.000001* | 1.995 | 0.256 | 0.000389* |
| Subiculum | 5.121 | 0.000007* | 1.619 | 0.229 | 0.000873* |
| CA2-CA3 | 5.025 | 0.000009* | 1.589 | 0.288 | 0.000142* |

Notes: ↑ rows are sorted based on the *p* values for the t-test; \* *p* < 0.008333 Bonferroni corrected for multiple comparison; *d* represents Cohen's d effect size; MMSE: mini mental state examination; SR-SL-SM: strata radiatum/lacunosum/moleculare

To investigate the relationship between the three whole-brain segmentation methods CAT, volBrain and BrainSuite, we ran correlational analysis, Table 6. Seven brain areas were common between these methods: nucleus accumbens, amygdala, caudate, globus pallidus, hippocampus, putamen and thalamus. CAT and volBrain showed strong correlation for nucleus accumbens, amygdala, caudate, hippocampus and thalamus. Two brain areas globus pallidus and putamen were not significantly correlated. These brain areas did not show significant difference between the two groups either. BrainSuite, however, showed no significant correlation with either of the other two segmentation methods.

**Table 6.** Correlation of the size of common brain areas reported by the three segmentation methods

|  |  | CAT |  | volBrain |  |
| --- | --- | --- | --- | --- | --- |
| | Brain Area | $r^2$ | $p$ | $r^2$ | $p$ |
| BrainSuite | Accumbens | -0.227 | 0.159813 | -0.277 | 0.078996 |
|  | Amygdala | 0.240 | 0.136249 | 0.029 | 0.858460 |
|  | Caudate | 0.169 | 0.298015 | 0.090 | 0.577545 |
|  | Globus Pallidus | -0.118 | 0.469153 | 0.184 | 0.249666 |
|  | Hippocampus | -0.162 | 0.318747 | -0.275 | 0.081327 |
|  | Putamen | 0.328 | 0.039077 | -0.186 | 0.243349 |
|  | Thalamus | 0.188 | 0.245249 | 0.177 | 0.268448 |
| volBrain | Accumbens | 0.633 | 0.000007* |  |  |
|  | Amygdala | 0.632 | 0.000007* |  |  |
|  | Caudate | 0.470 | 0.001678* |  |  |
|  | Globus Pallidus | -0.245 | 0.118543 |  |  |
|  | Hippocampus | 0.637 | 0.000006* |  |  |
|  | Putamen | 0.020 | 0.898315 |  |  |
|  | Thalamus | 0.541 | 0.000214* |  |  |

Notes: \* $p < 0.002380$ Bonferroni corrected for multiple comparison

## 4 Discussion

We used HIPS automated method to segment the subfields of hippocampus, and CAT, volBrain and BrainSuite automated methods to segment the whole brain using T1 weighted MRI data. Our results showed that all subfields of hippocampus in the Alzheimer’s Disease (AD) group were significantly smaller than those of the healthy control (HC) group. The atrophy of all subcomponents of hippocampus were correlated with the MMSE measure. Quite a large portion of cortical and subcortical areas in the brain were also smaller in the AD group as compared to the control group, as evident from CAT and volBrain segmentation results. The shrinkage in these brain areas mostly showed a strong correlation with MMSE measure. BrainSuite failed to discriminate between the two groups. While CAT and volBrain shows a strong correlation, BrainSuite did not show any significant correlation with CAT and volBrain.

With the advancement of computational methods, fine-grain analysis of the brain areas is more feasible ^73–75^. Earlier methods relied heavily on manual segmentation of the brain areas, which was extremely time demanding and also required a great level of expertise. Therefore, the majority of the analysis was limited to brain areas with more distinct structure, such as the hippocampus. Many semi- and fully-automated segmentation methods have been developed. While these methods have been used more commonly in recent years, the reliability and accuracy of these methods was yet to be fully studied. We used four pipelines of HIPS ^33^, volBrain ^34^, CAT ^35,36^ and BrainSuite ^37^ In this study we evaluated their reliability by looking at their ability to discriminate between AD and HC groups, whether a correlation existed between them, their correlation with MMSE scores, and comparing their results with past literature. Our results showed strong reliability of HIPS, volBrain and CAT. These methods have been successfully applied to brain images from those with AD ^76–79^

BrainSuite, however, underperformed greatly. For example, it failed to accurately segment the hippocampus, thalamus and amygdala to show a significant difference between the two groups. While this automatic segmentation method has been used frequently in past research ^80^, its application has been mostly limited to the processing of brains with no atrophy ^81,82^, as well as detection of gross segments such as tumours ^83^. This highlights the importance of validation studies such as ours to gain a greater understanding of the applications and limitations of different methods ^84–87^

The volume of the hippocampus is considered as an important biomarker for AD and has been included in recently proposed research diagnostic criteria ^88,89^. It has been shown that the hippocampal atrophy estimated on anatomical T1 weighted MRI can help in classifying the different stages of AD ^90–92^ Confirming past literature, our results showed that the hippocampus volume significantly differed between AD and the control group.

Histological studies have shown that lesions are not uniformly distributed within the hippocampus ^91,93^. Neuronal loss results in a reduction of the thickness of the layers richer in neuronal bodies, while the loss of synapses results in the reduction of the layers poorer in neuronal bodies ^94–97^ and these changes are stage-dependent ^98,99^ Our results, however, failed to differentiate the contribution of these subfields in AD; they all showed significant reduction in size, compared to the control group. This effect could be because our AD group consisted of those with later stages of AD. The contribution of different subfields of the hippocampus is more visible in those with mild cognitive impairment ^100–102^ Therefore, in future studies it would be informative to include participants with different stages of AD to investigate the contribution of different subfields of the hippocampus in AD.

While the contribution of atrophy in the hippocampus has been widely studied, the role of atrophy in the rest of the brain in AD is less clear ^40^. An important contributing factor is that the boundaries of the hippocampus are easier for human operators or automated algorithms to recognise than other brain areas such as the amygdala, entorhinal cortex or thalamus ^40^. Due to methodological advances, however, it is now possible to measure atrophy across the entire cortex with good precision ^103^. Our results from CAT and volBrain methods showed strongly significant differences between many brain areas such as the amygdala, thalamus, nucleus accumbens, insula and caudate. These findings are in-line with past literature showing similar differences in these brain areas ^40,104–106^.

There is a growing body of literature showing a correlation between cognitive decline and brain atrophy ^62,107,108^. For example, it has been shown that basal forebrain changes are correlated with cognitive decline in MCI and AD patients, as measured with recall task and MMSE ^109,110^, as well as healthy participants that later progressed to AD ^111^. Atrophy of other brain areas such as lateral and medial parietal cortex, as well as lateral temporal cortex have also been shown to have a correlation with cognitive decline ^112^ Our results showed a strong correlation between brain atrophy and cognitive decline as measured by MMSE. All brain areas that were significantly different between the AD and the control group showed a significant correlation with MMSE, except for the caudate (CAT *p* = 0.001155, volBrain *p* = 0.005091, Bonferroni corrected statistic not significant). While the effect of shrinkage of the caudate in AD is not very clear ^113^, there is some evidence that caudate volume has a correlation with MMSE measures, although not as strongly as other brain areas such as the thalamus ^63,114^ An important consideration is that atrophy in the left caudate has a stronger role in AD, as compared to the right caudate ^113,115^. Our analysis combined both the left and right caudate, which may have led to this inconsistency between our results and previous literature.

Although AD commonly presents as an amnestic syndrome, there is significant heterogeneity across individuals ^116^, which is accompanied by different atrophy patterns ^64,117–119^ For example, while those with more language difficulties might exhibit greater atrophy in temporal or parietal regions ^120,121^, those with more visual difficulties might have greater atrophy in posterior cortical regions ^122,123^. Availability of the automated systems offers many opportunities, such as the ability to analyse a large number of brain images with reasonable time and expertise. This is in particular very appealing, considering the increased number of large datasets such as MIRIAD. Automated systems can go through the collection and aggregate data from a wide range of participants, healthy and patients to gain a greater understanding of AD. This is important considering the heterogeneity of the disease and its progression ^124^.

Another application of automated systems is in clinical settings. By the time of diagnosis, rapid ongoing atrophy is already far advanced ^125,126^. Early diagnosis of AD can help with deceleration of the progression of the disease ^21,127,128^. This is particularly important as there are modifiable factors that can help with brain health^129–131^. Therefore, a massive effort has been devoted to the development of diagnostic methods to enable researchers and clinicians to detect AD and cases with potential progression to AD, as early as possible ^132,133^. For the development of preventive strategies, it is important to predict future brain atrophy, as this may aid in identifying which individuals with normal cognition are more susceptible of progressing to later stages of AD ^134–136^. Automated systems provide additional information to clinicians, enabling them to have a greater understanding of the progression of the atrophy ^12,137^ Some of these methods have already received approval from different licensing bodies such as CE (European conformity) and FDA (food and drug administration, USA) approval ^138^. These methods, however, come with some limitations such as speed of processing, expensive licences, or requirement of other specialised software. This study is another step to evaluate freely available analytical tools to achieve an ideal analysis pipeline, suitable for researchers and clinicians.

Availability of the reliable automated segmentation methods enables researchers and clinicians to have a greater understanding of the underlying mechanisms and the progression of the AD. This will allow them to attempt to prevent or decelerate the progression of the disease more effectively. Future research can look at the rate of atrophy to predict the progression of disease ^139–142^ This rate can be helpful to have a more informed understanding whether an individual with MCI will later progress to AD or not ^105,143,144^ The output of automated segmentation methods can also be used in training of intelligent classification methods such as those using artificial neural networks and support vector machines, which has shown promising results ^145–154^

The purpose of this article was not to identify the superiority of any particular automatic segmentation method over another, but to solely highlight possible limitations and applications of four commonly used segmentation methods. We proposed that CAT, volBrain and HIPS are methods that can robustly operate on brain images with significant atrophy and can be used in research and clinical settings. BrainSuite, however, should be used with caution for brain images with atrophy.

## Authors’ Contribution

JZ and AHJ analysed the data. JZ and AHJ wrote the manuscript. JZ, AHJ and AS revised the manuscript.

