## Supplementary for "A Large-scale Comparison of Cortical and Subcortical Structural Segmentation Methods in Alzheimer’s Disease: a Statistical Approach"

Supplementary Figure 15. Correlation of the common brain areas between volBrain and CAT. Red and blue dots indicate AD and HC respectively. .. 17

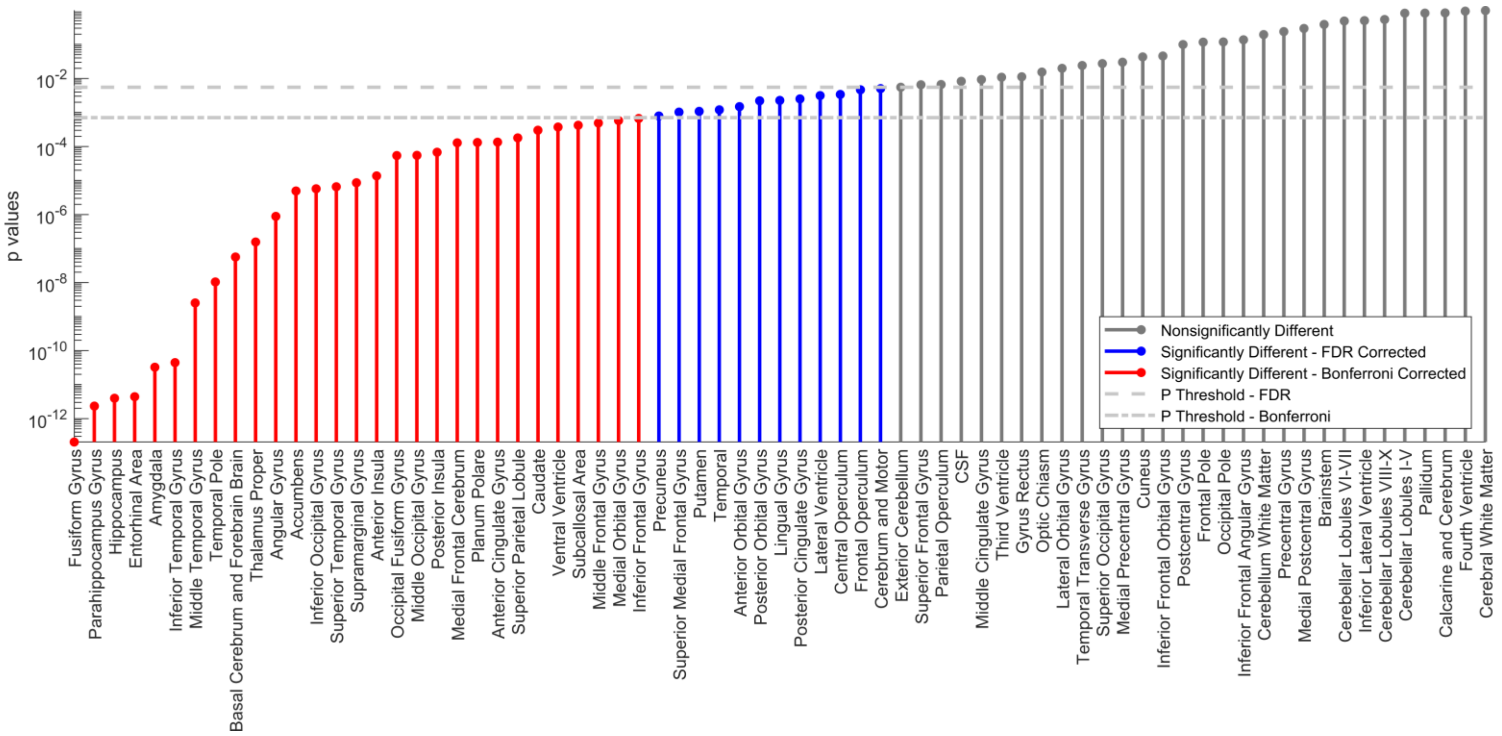

Supplementary Figure 1. Independent sample t-tests, comparing the volume of different segments between AD and HC using CAT. Brain areas are sorted based on the p-values. False discovery rate (FDR) [1] and Bonferroni correction have been used to correct for multiple comparisons.

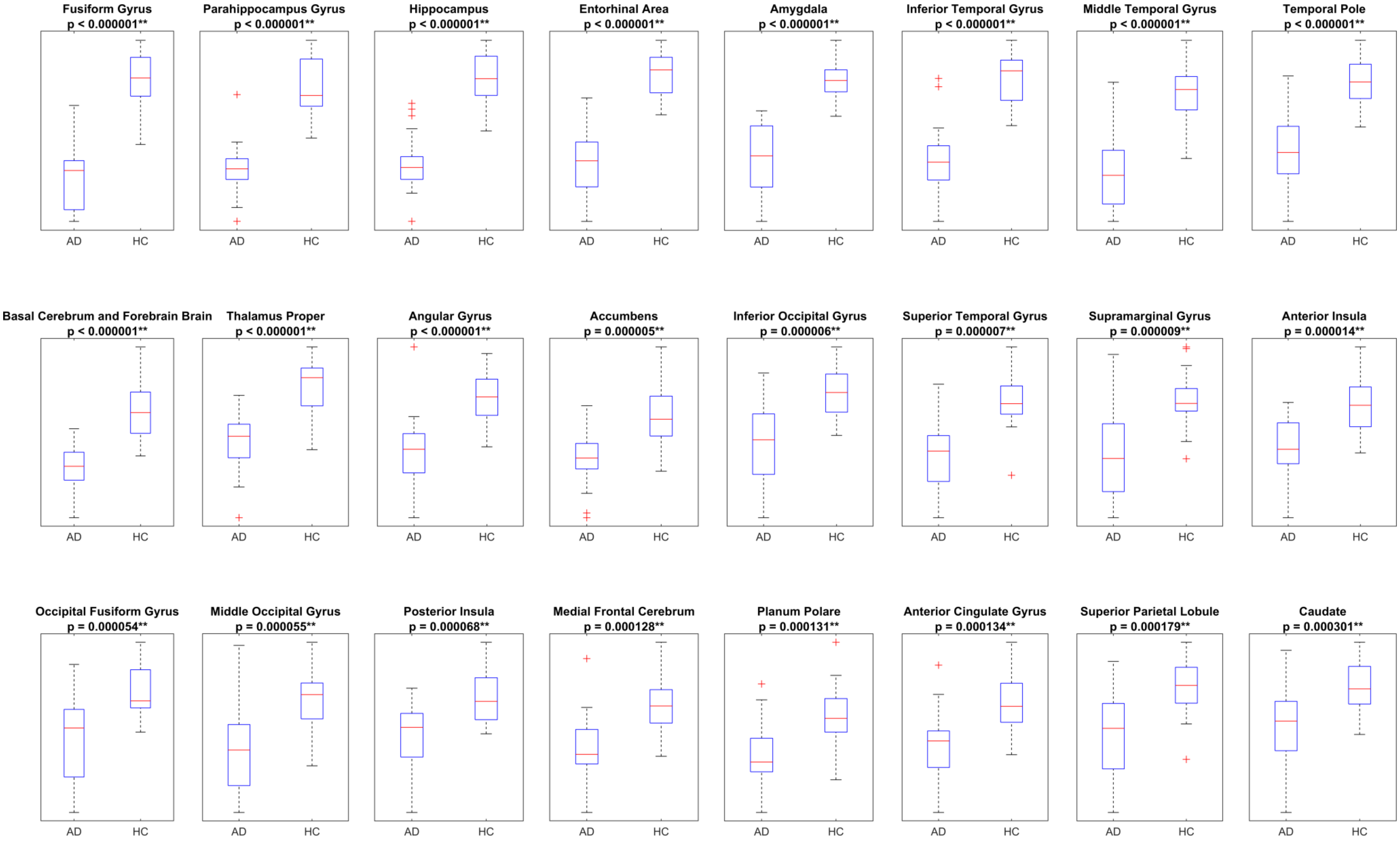

Supplementary Figure 2. Distribution of the volume of different segments in AD and HC using CAT, based on the first 24 brain areas indicated in Supplementary Figure 1

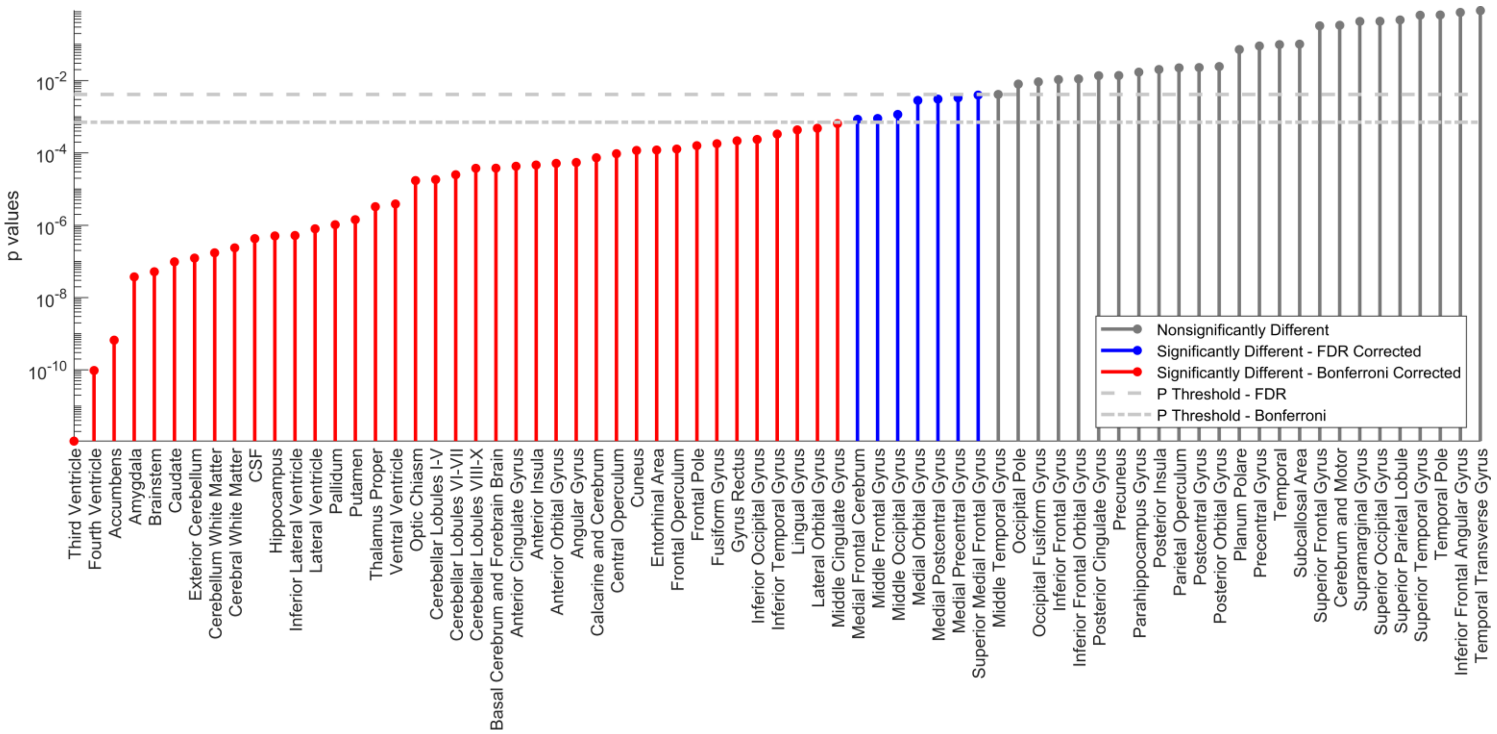

Supplementary Figure 3. Correlations of the volume of different segments with MMSE scores for both AD and HC using CAT. Brain areas are sorted based on the p-values. False discovery rate (FDR) [1] and Bonferroni correction have been used to correct for multiple comparisons.

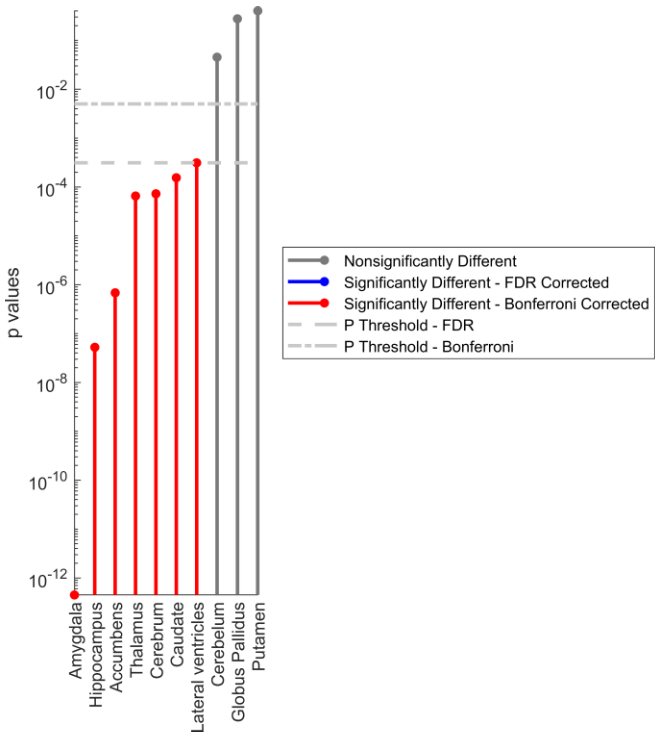

Supplementary Figure 4. Independent sample t-tests, comparing the volume of different segments between AD and HC using volBrain. Brain areas are sorted based on the p-values. False discovery rate (FDR) [1] and Bonferroni correction have been used to correct for multiple comparisons.

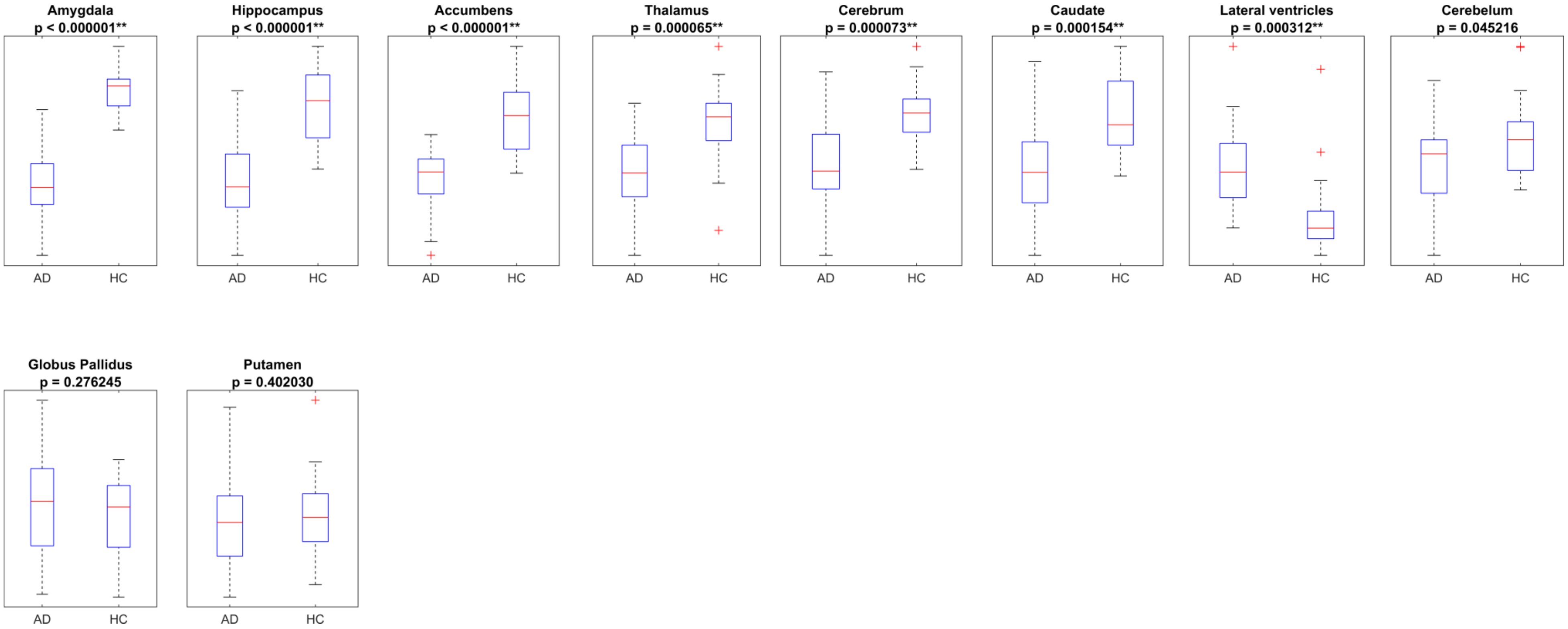

Supplementary Figure 5. Distribution of the volume of different segments in AD and HC using volBrain.

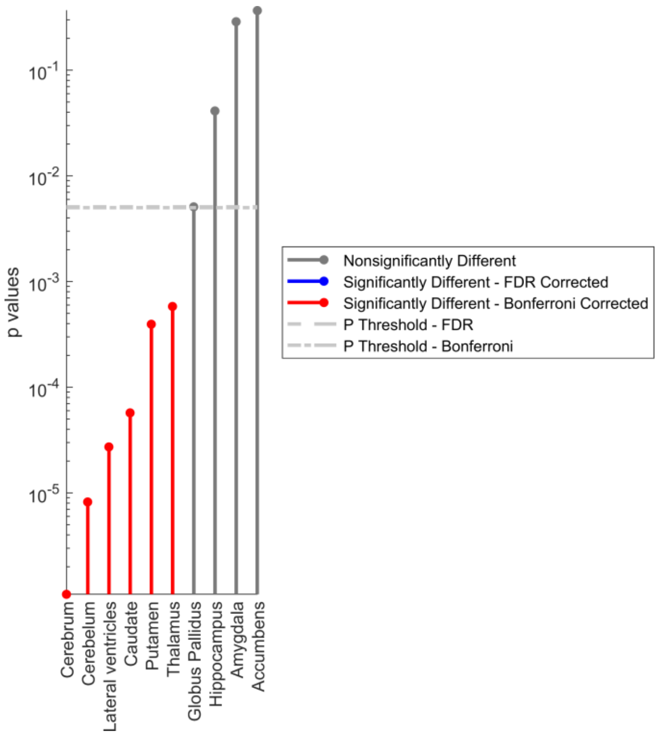

Supplementary Figure 6. Correlations of the volume of different segments with MMSE scores for both AD and HC using volBrain. Brain areas are sorted based on the p-values. False discovery rate (FDR) [1] and Bonferroni correction have been used to correct for multiple comparisons.

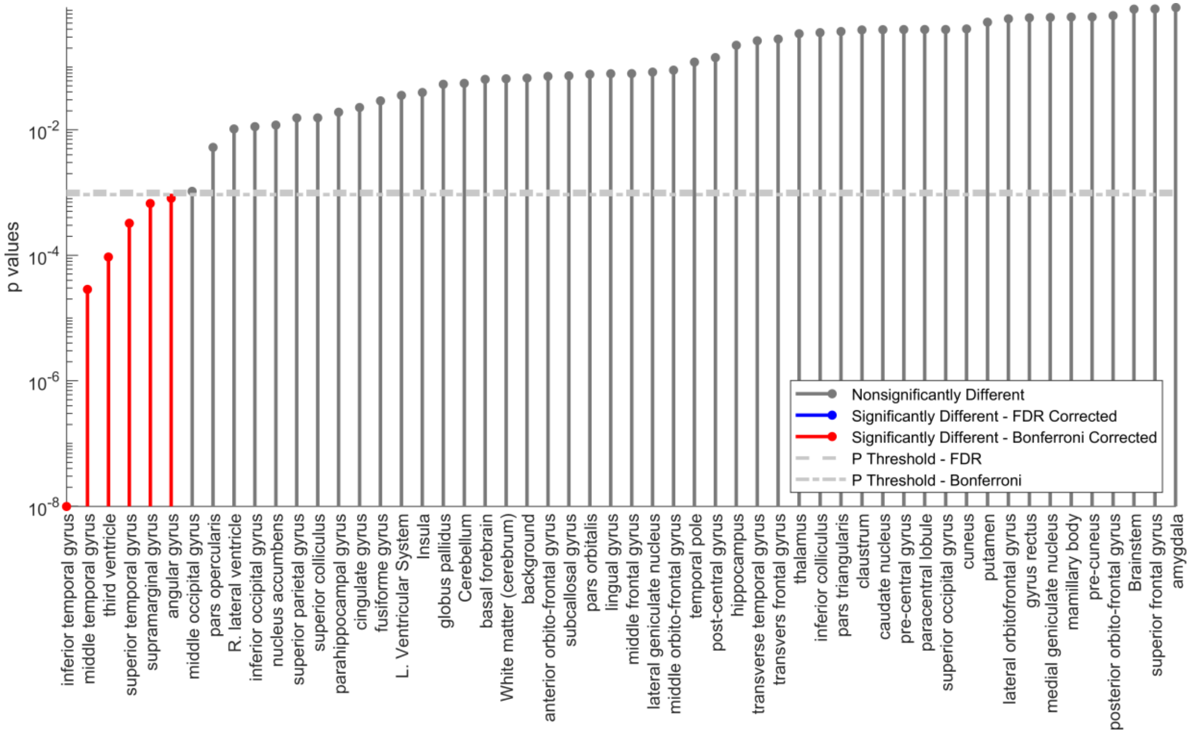

Supplementary Figure 7. Independent sample t-tests, comparing the volume of different segments between AD and HC using BrainSuite. Brain areas are sorted based on the p-values. False discovery rate (FDR) [1] and Bonferroni correction have been used to correct for multiple comparisons.

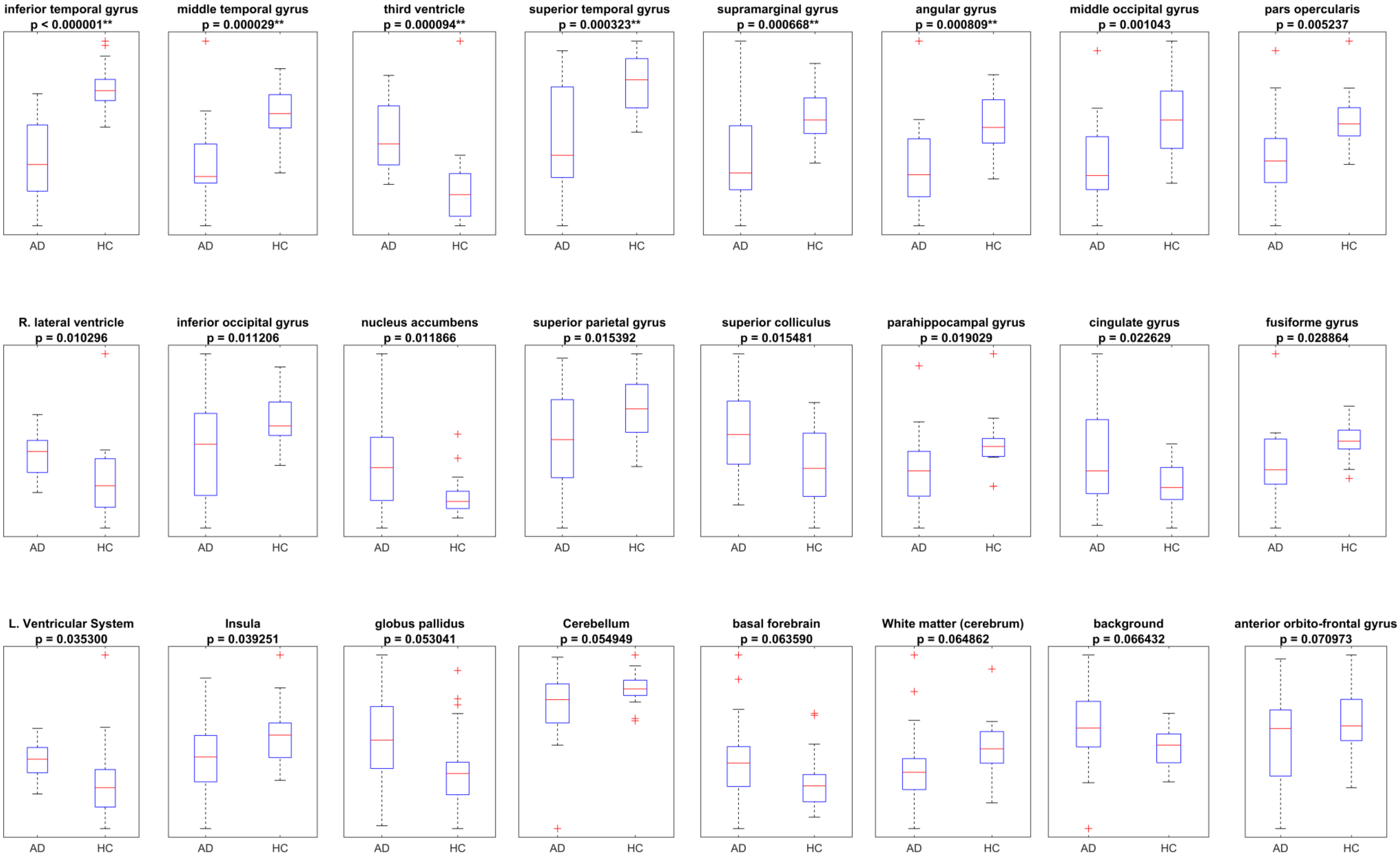

Supplementary Figure 8. Distribution of the volume of different segments in AD and HC using BrainSuite, based on the first 24 brain areas indicated in Supplementary Figure 7.

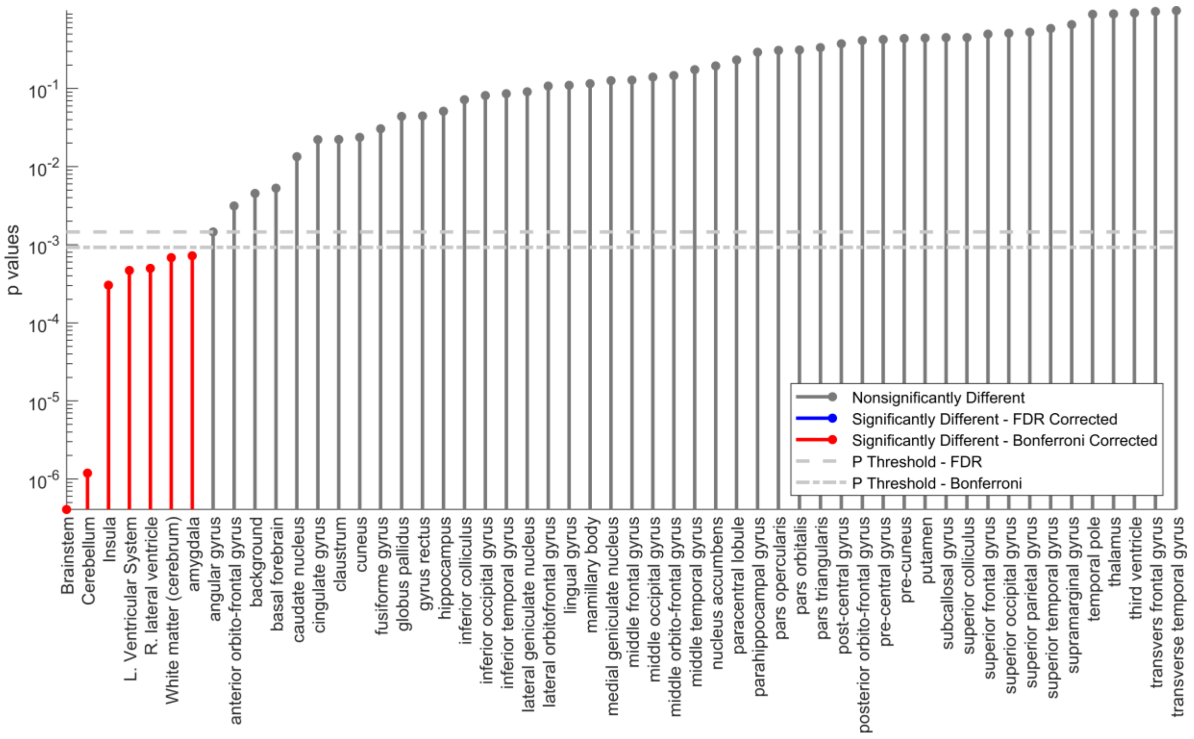

Supplementary Figure 9. Correlations of the volume of different segments with MMSE scores for both AD and HC using BrainSuite. Brain areas are sorted based on the p-values. False discovery rate (FDR) [1] and Bonferroni correction have been used to correct for multiple comparisons.

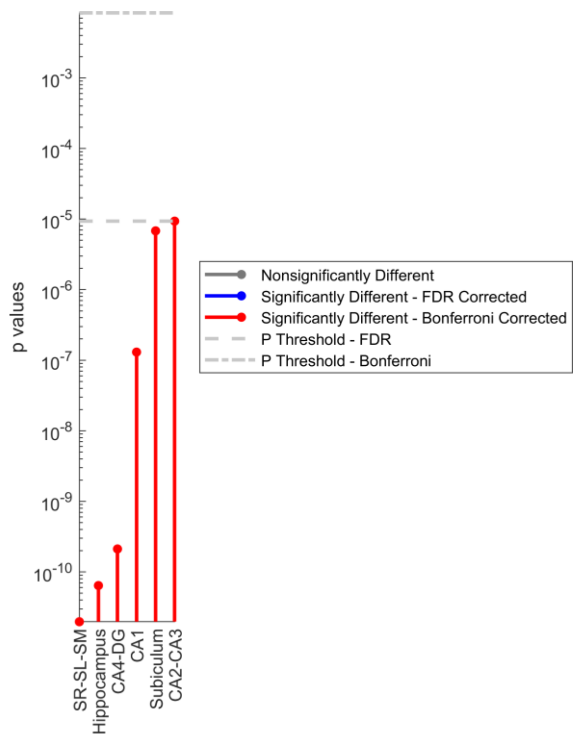

Supplementary Figure 10. Independent sample t-tests, comparing the volume of different subfields of hippocampus between AD and HC using HIPS. Brain areas are sorted based on the p-values. False discovery rate (FDR) [1] and Bonferroni correction have been used to correct for multiple comparisons.

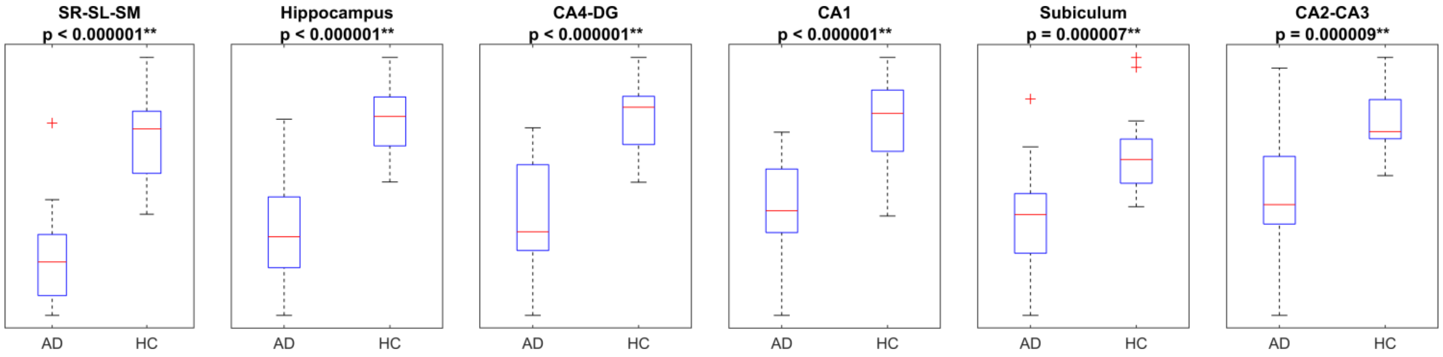

Supplementary Figure 11. Distribution of the volume of different subfields of hippocampus in AD and HC using HIPS.

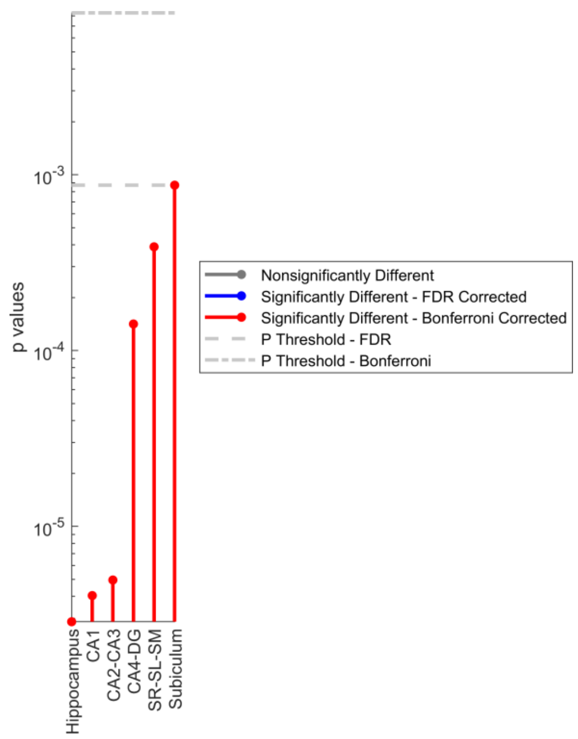

Supplementary Figure 12. Correlation of the volume of different subfields of hippocampus with MMSE scores for both AD and HC using HIPS. Brain areas are sorted based on the p-values. False discovery rate (FDR) [1] and Bonferroni correction have been used to correct for multiple comparisons.

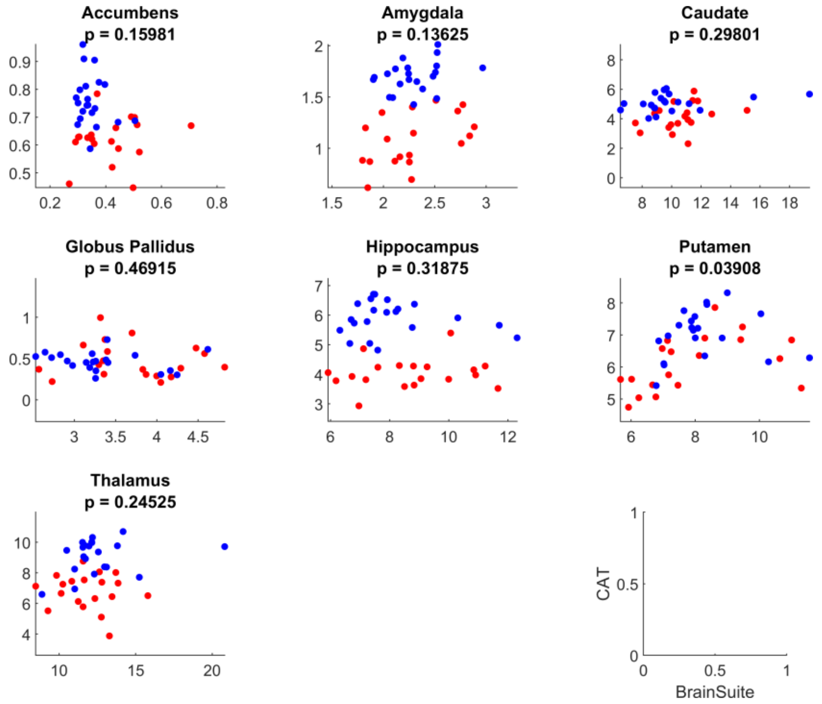

Supplementary Figure 13. Correlation of the common brain areas between BrainSuite and CAT. Red and blue dots indicate AD and HC respectively.

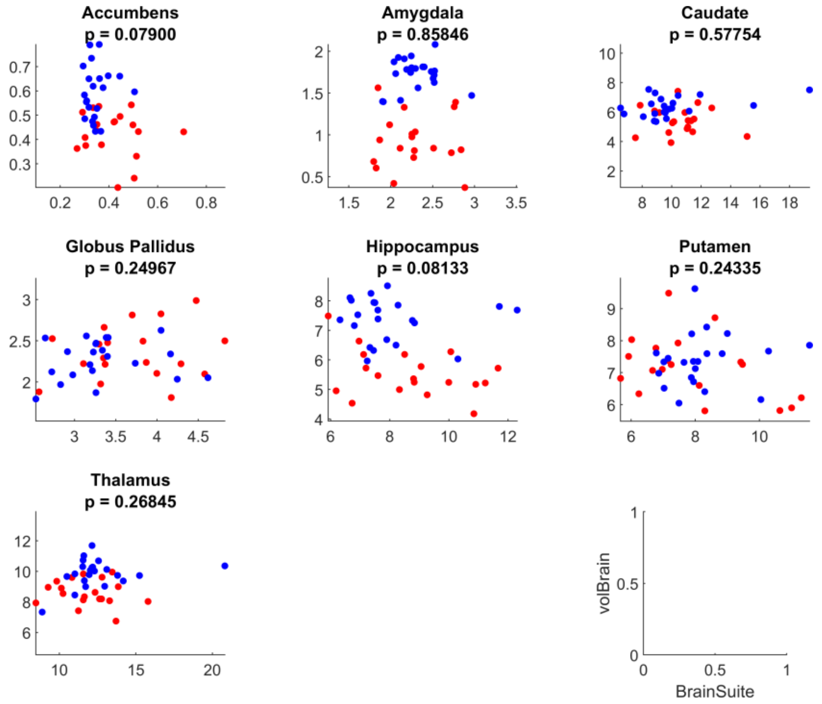

Supplementary Figure 14. Correlation of the common brain areas between volBrain and BrainSuite. Red and blue dots indicate AD and HC respectively.

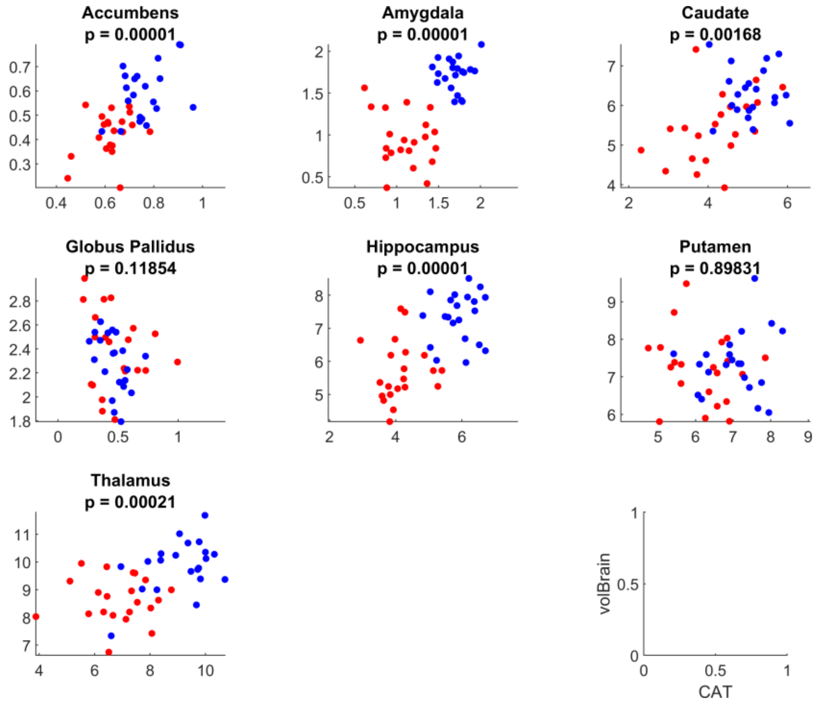

Supplementary Figure 15. Correlation of the common brain areas between volBrain and CAT. Red and blue dots indicate AD and HC respectively.
